# Microbial succession in casing layer shapes bacterial blotch disease associated communities in cultivated white button mushroom: From casing layer to disease

**DOI:** 10.64898/2026.09.10.750680

**Authors:** Sameerika D Mudiyanselage, Romina Gazis, Samuel J Martins

## Abstract

**Background:** Bacterial blotch is a major disease of cultivated white button mushroom (*Agaricus bisporus*) traditionally attributed to individual *Pseudomonas* pathogens. However, the recurrent detection of diverse bacterial taxa in blotch-affected mushrooms suggests that disease may involve broader changes in microbial community organization. We characterized bacterial communities associated with symptomatic and asymptomatic mushrooms and examined bacterial succession in the casing layer across early, pinning, and harvest stages at two commercial mushroom farms in the United States using complementary 16S rRNA gene amplicon sequencing and shotgun metagenomics.

**Results:** Mushroom-associated communities were dominated by *Pseudomonas* regardless of disease status, indicating that bacterial blotch was not simply associated with increased abundance of the dominant genus. Instead, symptomatic mushrooms exhibited significant community restructuring, enrichment of specific taxa, and markedly reduced microbial network complexity. Species-level metagenomics revealed extensive reorganization within *Pseudomonas*, with contrasting shifts among multiple blotch-associated lineages, accompanied by changes in non-*Pseudomonas* taxa, including *Mycetocola* and *Ewingella*. Despite these taxonomic shifts, dominant *Pseudomonas* populations retained broadly conserved functional profiles, with disease-associated enrichment of pathways related to central metabolism, O-antigen biosynthesis, and peptidoglycan maturation. Casing communities underwent pronounced directional succession, shifting from early dominance by *Exiguobacterium* and *Leuconostoc* toward enrichment of *Flavobacterium*, *Pedobacter*, and *Pseudomonas* at later stages. Multiple blotch-associated *Pseudomonas* lineages were detected throughout casing development, while *Pseudomonas* increased from approximately 2% in early casing to 51% at harvest at the farm with higher disease incidence. Succession occurred without significant changes in alpha diversity, indicating that community development primarily reflected taxon replacement and redistribution.

**Conclusions:** Our findings support a microbiome-centered framework for bacterial blotch in which disease is associated with host and stage dependent microbial succession, species-level community restructuring, and altered microbial connectivity. This framework extends beyond the single-pathogen paradigm and highlights bacterial blotch as a community-level disease process shaped by dynamic interactions between the mushroom host and its surrounding microbiome.

## Background

The white button mushroom (*Agaricus bisporus*) is the most widely cultivated edible fungi worldwide and is valued for its nutritional and functional properties, including dietary fiber, protein, essential minerals, unsaturated fatty acids, and a diverse array of bioactive compounds [1,2,3]. Beyond its economic and nutritional importance, the mushroom fruiting body represents a unique microbial habitat. Its high moisture content, favorable surface pH, and lack of protective cuticle create ecological niches that support diverse bacterial communities capable of colonizing fruiting tissues [4]. These microbial associations can influence mushroom growth, development, and productivity, but they may also contribute to disease emergence [5]. Consequently, understanding how microbial communities assemble and interact on mushroom surfaces has become increasingly important for sustainable mushroom production.

Bacterial blotch is among the most economically important diseases affecting cultivated *A. bisporus.* The disease is characterized by yellow to brown discoloration of the mushroom surface that may progress to pitting, tissue collapse, and substantial losses in market quality. Historically, bacterial blotch has been attributed primarily to *Pseudomonas tolaasii* and *P. “gingeri”* [6,7]. However, this pathogen centered framework has become increasingly difficult to reconcile with the growing diversity of bacteria recovered from blotch-affected mushrooms. Recent studies have identified approximately 20 *Pseudomonas* species associated with blotch symptoms, while bacteria belonging to other genera, including *Mycetocola*, *Serratia*, *Pantoea* and *Cedecea*, have also been isolated from symptomatic tissues [5,8,9,10,11,12]. The repeated detection of diverse bacterial taxa raises a fundamental ecological question: does bacterial blotch result from the activity of a single pathogen, or does disease emerge from a particular microbial community state in which multiple bacterial populations, interactions, and functional capacities collectively contribute to symptom development?

Addressing this question requires moving beyond traditional culture-based approaches. Although isolation-based studies have been instrumental in identifying bacterial taxa associated with blotch disease, they capture only the subset of microorganisms present in mushroom-associated communities. Culture-independent approaches, including 16S rRNA gene amplicon sequencing and shotgun metagenomics, enable comprehensive characterization of microbial communities while simultaneously providing insights into their potential functional capabilities. These approaches allow disease-associated populations to be studied within their broader ecological context rather than as isolated taxa. Despite growing recognition of the importance of microbiomes in crop health, the community-level organization of the *A. bisporus* bacteriome during bacterial blotch development remains poorly understood. This knowledge gap is particularly important because disease symptoms are expressed directly on the mushroom cap, where microbial interactions likely play a central role in determining disease outcomes. Previous studies have demonstrated that mushrooms caps harbor reproducible bacterial assemblages dominated by members of the *Proteobacteria* and *Bacteroidetes,* suggesting that disease may arise from perturbations within an established microbial community rather than from the introduction of a single pathogenic species [13].

The origin and temporal dynamics of these disease-associated bacterial communities may be closely linked to the microbial ecology of the mushroom production environment. During commercial cultivation, mushrooms development occurs within a series of interconnected microbial habitats, including compost and casing material, that undergo continuous biological and physiochemical change. Among these habitats, the casing layer is particularly important because it forms the immediate ecological interface between the colonized substrate and developing fruiting bodies. In addition to providing the physical and environmental conditions required for mushroom production, the casing serves as a reservoir of microorganisms capable of influencing the mushroom growth and development [14]. Certain casing-associated bacteria, such as *Pseudomonas putida,* have been implicated in the degradation of self-inhibitory volatile compounds produced during mycelial growth, thereby promoting fruit-body formation of the mushroom [15] At the same time, the casing layer has also been suspected to serve as a potential source of bacteria associated with blotch disease [7].

Importantly, casing-associated bacterial communities are not static. Instead, they undergo predictable successional changes in response to the mushroom development, shifts in resource availability, and changing environmental conditions throughout crop production [16,17]. Such succession can alter the composition and abundance of bacterial populations available to colonize emerging fruiting bodies. Consequently, temporal dynamics within the casing microbiome may influence the assembly of mushroom-associated bacterial communities and create ecological conditions that favor or suppress disease-associated populations. A key unresolved question is therefore whether bacteria associated with blotch disease are embedded within a temporally structured casing microbiome that precedes disease development and may serve as a reservoir for disease-associated populations.

In a recent study, we characterized bacterial communities associated with bacterial blotch in cultivated white button mushrooms collected from two commercial farms in the United States and identified more than 17 bacterial species associated with symptomatic fruiting bodies [5]. Building upon those findings, we investigated bacterial community assembly across both mushroom cap tissues and casing material within the same production systems. We combined 16S rRNA gene amplicon sequencing and shotgun metagenomics to characterize the taxonomic composition and functional potential of bacterial communities associated with symptomatic and asymptomatic mushrooms while simultaneously tracking bacterial succession within the casing layer across early, pinning and harvest stages of mushrooms production. We hypothesized that bacterial blotch is associated with distinct microbiome states and that disease-associated bacterial populations are embedded within the successional dynamics of casing-associated microbial communities before disease symptoms become apparent. By linking disease-associated bacterial communities in mushroom tissues with microbial succession in the casing environment, this study provides an ecological framework for understanding bacterial blotch as a microbiome-associated disease rather than solely a pathogen-driven process. Specifically, our objectives were to determine whether (i) symptomatic and asymptomatic mushrooms harbor distinct bacterial community states, (ii) specific bacterial taxa and functional capacities are consistently associated with disease status (iii) casing-associated bacterial communities undergo directional succession throughout mushrooms development, and (iv) bacterial blotch-associated populations are detectable within casing communities prior to fruit-body formation and change in abundance over time.

## Material and Methods

### Bacterial blotch-associated mushroom and casing sampling

White button mushrooms (*Agaricus bisporus*) were collected from two commercial mushroom farms in the United States between April and May 2025, hereafter referred to as Farm X and Farm Y. A total of 60 mushrooms were sampled, including 30 mushrooms exhibiting bacterial blotch symptoms (symptomatic) and 30 asymptomatic mushrooms. Symptomatic and asymptomatic mushrooms were collected as paired samples approximately 50 cm apart to minimize environmental variation while reducing the risk of cross-contamination. At Farm X, 17 symptomatic and 17 asymptomatic mushrooms were collected, whereas 13 symptomatic and 13 asymptomatic mushrooms were collected from Farm Y.

Casing material was collected concurrently from the same farms representing at three developmental stages: (i) early-stage casing immediately after application and pasteurization, (ii) casing during mushroom pinning, and (iii) casing at harvest. Approximately 10 g of casing material was collected at a depth of 5 cm from randomly selected locations within mushroom beds. Three replicate samples were collected from each developmental stage at each farm, resulting in a total of 18 casing samples. Individual mushrooms and casing samples were placed into sterile plastic bags, transported to the laboratory under refrigerated conditions within 24 to 48 h, and stored at - 80°C until DNA extraction and microbiome analyses.

### DNA extraction, 16S rRNA amplicon sequencing, and shotgun metagenomic sequencing

Individual mushroom samples were aseptically transferred to sterile stomacher bags containing 5 mL of 10 mM phosphate buffer (pH 7.0) and macerated on ice. A 250 µL aliquot of the resulting suspension was used for Genomic DNA (gDNA) extraction. For casing samples, 200 mg of material was used for genomic DNA extraction. Genomic DNA was extracted from all mushroom samples (*n* = 60) and casing samples (*n* = 18) using the Quick-DNA Fecal/Soil Microbe MiniPrep Kit (Zymo Research, Irvine, CA, USA) according to the manufacturer’s instructions. For shotgun metagenomic sequencing, representative samples were selected from Farm X because this farm exhibited higher bacterial blotch incidence than Farm Y. Selected samples included casing collected at three developmental stages (*n* = 9) and symptomatic and asymptomatic mushroom samples (*n* = 6 each).

DNA quality was assessed by agarose gel electrophoresis using a Bio-Rad Gel Doc XR imaging system, and DNA concentrations were measured using a NanoDrop spectrophotometer (Thermo Fisher Scientific, Waltham, MA, USA). DNA samples were submitted to SeqCenter (Pittsburgh, PA, USA) for both 16S rRNA amplicon sequencing and shotgun metagenomic sequencing.

### 16S rRNA gene amplicon sequencing and bioinformatic analyses

Bacterial communities were characterized by amplification and sequencing of the V3-V4 region of the 16S rRNA gene using the universal primer pair 341F and 806R. Following library preparation, cleanup, and normalization, amplicons were sequenced on an Illumina NextSeq 2000 platform using a P1 or P2 600-cycle flow cell to generate paired-end reads. Sequence processing was performed using QIIME 2 (v2025.7) [18]. Raw sequence quality profile was inspected prior to analysis, and primer sequences were removed using cutadapt plugin implemented in QIIME 2. Denoising, quality filtering, chimera removal, and amplicon sequence variant (ASV) inference were performed using DADA2 [19]. Based on sequence quality profiles, forward and reverse reads were truncated at 270 bp and 220 bp, respectively. The resulting ASV feature table, and representative sequences were used for downstream analyses.

Taxonomic classification was performed using the q-2 feature-classifier plugin [20] with a pre-trained naïve Bayes classifier based on the SILVA 138 reference database clustered at 99% sequence identity [21]. Sequences identified as chloroplasts or mitochondria were removed prior to downstream analyses. Phylogenetic relationships among ASVs were inferred using the MAFFT [22] and FastTree [23] workflow implemented in QIIME 2. Representative sequences were aligned with MAFFT, hypervariable regions were masked, and rooted and unrooted phylogenetic trees were generated. Rarefaction analysis was performed using the rooted phylogenetic tree and ASV table to evaluate whether sequencing depth was sufficient to capture bacterial diversity, using a sampling depth of 70,000 sequences per sample. Taxonomic composition was summarized as relative abundance and visualized in R (v4.6.0). Core bacterial taxa were identified using the q-2 core-features approach, with core taxa defined as those present in at least 50% of samples. Shared and unique core taxa among sample groups were visualized using the Ven diagram. Alpha diversity was evaluated using Faith’s phylogenetic diversity [24], Shannon diversity [25], and Pielou’s evenness [26] Differences among mushroom health states (symptomatic and asymptomatic) and casing developmental stages (early, pinning, and harvest) and were assessed using Kruskal-Wallis tests [27] followed by pairwise Wilcoxon rank-sum tests [28] when appropriate. Beta diversity was assessed using Bray-Curtis [29], generated within the q-2 core diversity pipeline. Principal coordinates analysis (PCoA) was used to visualize community dissimilarities. Differences in community composition among sample groups were evaluated using permutational multivariate analysis of variance (PERMANOVA) with 999 permutations [30]. Pairwise PERMANOVA comparisons were conducted where appropriate, and *p*-values were adjusted using the Benjamini– Hochberg false discovery rate (FDR) correction procedure [31]. Significant was determined at *q* < 0.05.

Differential abundance analysis was performed using ANCOM-BC [32] implemented in QIIME 2. Taxa represented by fewer than 50 total reads across all samples were removed prior to analysis. Differentially abundant genera associated with mushroom health status or casing developmental stage were identified using FDR-adjusted significance thresholds (*q* < 0.05). Results were exported to R for visualization, and genera were ranked according to log2 fold change. For presentation, the 15 most enriched and 15 most depleted genera were displayed, with *Pseudomonas* retained regardless of ranking because of its biological relevance to bacterial blotch disease.

### Co-occurrence network analysis

Co-occurrence networks were constructed separately for asymptomatic and symptomatic mushroom microbiomes. To minimize the influence of rare taxa, analyses were restricted to genera detected in at least 20% of samples within each health-status group. Relative abundance tables were normalized to the total microbial abundance of each sample prior to network construction. Pairwise associations among bacterial genera were calculated using Spearman’s rank correlation coefficients implemented in the R package Hmisc. Significant associations were retained when the absolute correlation coefficient exceeded 0.60 ((|ρ|) ≥ 0.6) and the associated significance value was *p* < 0.001. Non-significant and weak associations were removed. Undirected weighted networks were generated using the R package igraph. Positive and negative correlations were represented as separate edge types. Network topology was characterized using the number of nodes, number of edges, network density, average degree, clustering coefficient, and modularity. Network visualization was performed using a Fruchterman-Reingold force-directed layout, with node size scaled according to mean relative abundance of each bacterial genus.

### Shotgun metagenomic sequencing and bioinformatic analysis

Illumina sequencing libraries were prepared using the tagmentation-based Illumina DNA Prep kit with custom Integrated DNA Technologies (IDT) 10 bp unique dual indices (UDI), targeting an average insert size of approximately 280 bp. No additional DNA fragmentation or size-selection steps were performed. Libraries were sequenced on an Illumina NovaSeq X Plus platform in multiplexed runs, generating 2 × 151 bp paired end reads. Primary demultiplexing, adapter trimming, and initial quality filtering were performed using bcl-convert (v4.2.4). Raw sequencing quality was assessed using FastQC (v0.12.1). Additional read processing was performed using Trim Galore (v0.6.10), where reads were quality-filtered using a Phred score threshold of 20. The first 15 bases from the 5′ end and the last 10 bases from the 3′ end of both forward and reverse reads were trimmed to remove low-quality and biased sequence regions. High-quality paired-end reads were retained for downstream analyses. Taxonomic profiling was performed using Kraken2 (v2.1.3) [33] with the PlusPF database, which includes bacterial, archaeal, viral, fungal, protozoan, plant, and vector sequences derived from RefSeq and UniVec_Core. Species-level abundance estimates were refined using Bracken (v2.7.0) [34], which recalculates taxon abundances using genome-specific k-mer distributions and read length information. Taxonomic profiles were used for downstream statistical and compositional analyses.

Microbial functional potential was characterized using HUMAnN v3.8. Pathway abundance profiles generated by HUMAnN were used for downstream analyses of metabolic potential and community functional composition. Temporal changes in bacterial and *Pseudomonas* assemblages across casing developmental stages were visualized using alluvial plots generated with the ggalluvial package in R. The relative abundances of bacterial blotch-associated *Pseudomonas* species, including *P. tolaasii*, *P. gingeri*, *P. agarici*, *P. reactans*, *P. yamanorum*, *Pseudomonas* sp. NC02, *P. azotoformans*, *P. pergaminensis*, *P. monsensis*, *P. tensinigenes*, *P. edaphica*, *P. salomonii*, *P. extremorientalis*, *P. simiae*, *Pseudomonas* sp. Irchel 3A7, and *Pseudomonas* sp. REP124, were visualized using heatmaps generated in ggplot2 in R.

## Results

### Bacterial blotch is associated restructuring of the white button mushroom bacteriome

Sixty white button mushrooms collected from two commercial farms were analyzed, including symptomatic mushrooms expressing bacterial blotch symptoms and asymptomatic mushrooms without visible symptoms. Symptomatic mushrooms exhibited both ginger blotch, characterized by ginger-colored discoloration, cap sliminess, and surface cracking, and brown blotch, characterized by sunken brown lesions on mushrooms caps (Fig. 1A). Amplicon sequencing generated 2,737 bacterial amplicon sequence variants (ASVs), with an average of 74,450 reads per sample (Supplementary Excel File 1).

**Figure 1.**
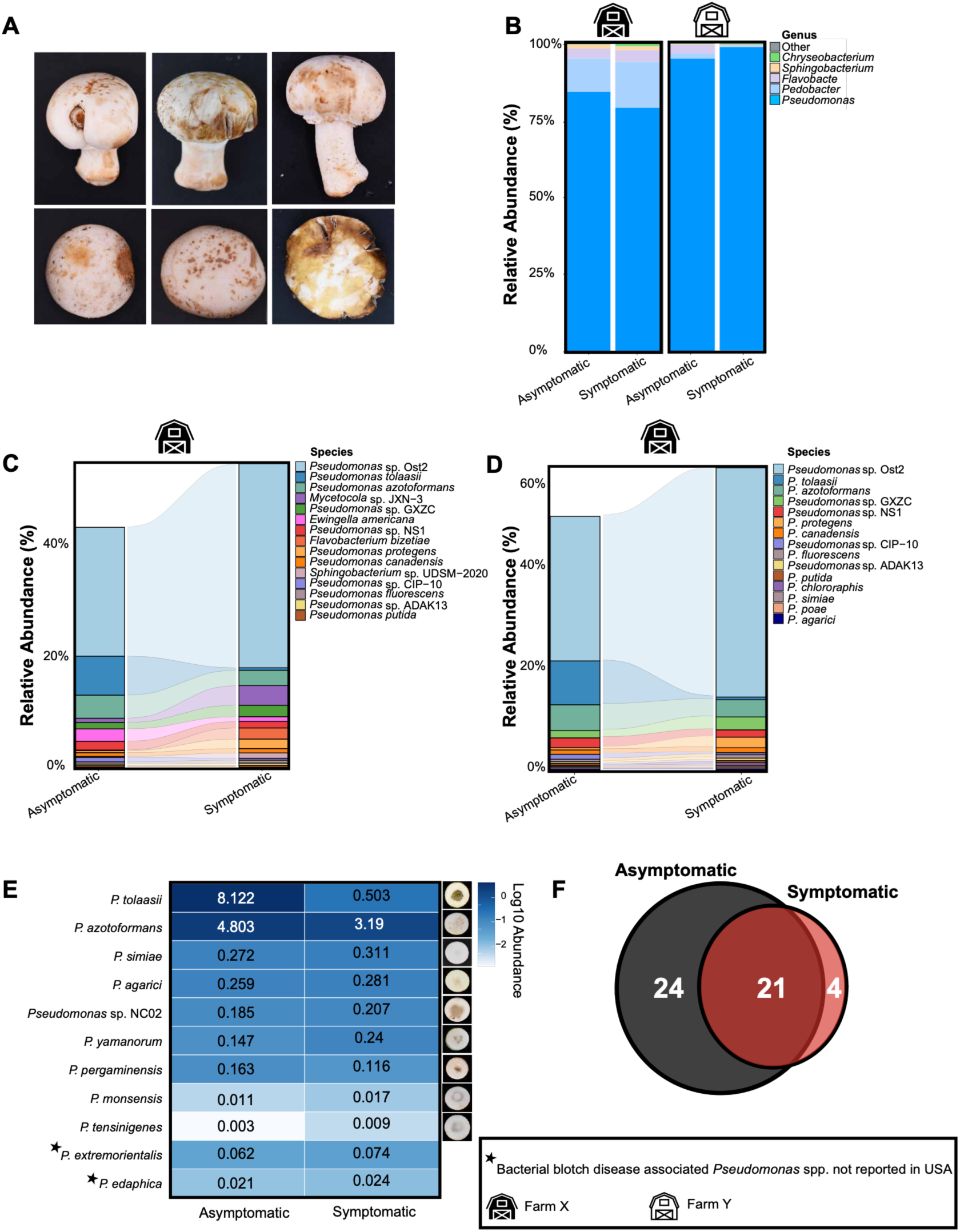
Bacterial blotch-associated bacterial communities in white button mushroom. (A) Bacterial blotch symptoms on white button mushrooms, ranging from dark brown lesions to ginger colored discoloration, accompanied by varying degrees of tissue pitting. (B) Genus-level relative abundance of the top five bacterial genera associated with asymptomatic and symptomatic mushrooms from Farm X and Farm Y. The category “Other” represents all remaining bacterial genera. (C) Species-level relative abundance of the top 15 bacterial taxa in asymptomatic and symptomatic mushrooms from Farm X. (D) Relative abundance of the top 15 *Pseudomonas* species detected in asymptomatic and symptomatic mushrooms from Farm X. (E) Heatmap showing the relative abundance of bacterial blotch associated pathogenic *Pseudomonas* species in asymptomatic and symptomatic mushrooms from Farm X. Representative symptoms associated with each *Pseudomonas* species are shown in the right panel based on symptoms observed for isolates previously characterized by Mudiyanselage et al. (2026). Values represent mean Log_10_-transformed abundance. (F) Venn diagram showing the numbers of bacterial ASVs shared among and exclusively detected at the symptomatic mushrooms and asymptomatic mushrooms. * indicate *Pseudomonas* species associated with bacterial blotch disease that have not previously been reported in the United States. Farm labels indicate the sampling locations (Farm X and Farm Y).

Genus-level community profiling revealed that the mushroom-associated bacterial communities were dominated by *Pseudomonas*, *Pedobacter*, *Flavobacterium*, *Sphingobacterium*, and *Chryseobacterium* (Fig. 1B and Supplementary Excel File 1). *Pseudomonas* was the predominant genus regardless of disease status. In Farm X, *Pseudomonas* represented approximately 78% and 77% of the bacterial community in asymptomatic and symptomatic mushrooms, respectively. In Farm Y, the relative abundance of *Pseudomonas* increased from 87% in asymptomatic mushrooms to 97% in symptomatic mushrooms. Despite these patterns, both symptomatic and asymptomatic mushrooms remained strongly dominated by *Pseudomonas*-associated communities.

To obtain species-level taxonomic resolution, six mushroom samples from Farm X, which exhibited the higher disease incidence, were subjected to shotgun metagenomic sequencing. Metagenomic sequencing generated 77.5 million paired-end reads, of which 74.6 million high-quality reads were retained after quality filtering. Individual samples yielded between 8.1 to 16.1 million reads, with an average GC content of 51%, indicating consistent sequencing quality and genomic coverage across samples (Supplementary Excel File 1).

Species-level profiling revealed substantial restructuring of bacterial communities associated with disease status (Fig. 1C and Supplementary Excel File 1). *Pseudomonas* sp. Ost2 was the dominant species and increased from approximately 26% in asymptomatic mushrooms to 42% in symptomatic mushrooms. In contrast, *P. tolaasii*, *P. azotoformans*, and *Ewingella americana* decreased in symptomatic mushrooms. Other taxa, including *Mycetocola* sp. and *Pseudomonas* sp. GXZC, were enriched in symptomatic samples. Because *Pseudomonas* dominated all mushroom-associated bacterial communities, species-level dynamics within this genus were further examined (Fig. 1D). Multiple *Pseudomonas* taxa differed in relative abundance between symptomatic and asymptomatic mushrooms. Some blotch-associated species declined in symptomatic mushrooms, whereas others increased, indicating substantial restructuring within the *Pseudomonas* assemblage rather than a uniform increase across all members of the genus.

Screening metagenomic datasets against previously reported bacterial blotch-associated *Pseudomonas* species identified eleven taxa in commercial mushrooms production systems in the United States (Fig. 1E). These include *P. tolaasii, P. azotoformans, P. simiae, P. agarici, P. yamanorum, P. tensinigenes, P. monsensis, P. pergaminensis* and *Pseudomonas* sp. NC02. Relative abundances varied substantially between symptomatic and asymptomatic mushrooms. In addition, *P. extremorientalis* and *P. edaphica,* previously associated with bacterial blotch outside the United States, were also detected.

### Symptomatic and asymptomatic mushrooms harbor distinct bacterial community states

Core microbiome analysis revealed broadly overlapping but differentially represented bacterial assemblages in symptomatic and asymptomatic mushrooms (Fig. 1F). Four unique core ASVs were detected in symptomatic mushrooms and 24 in asymptomatic mushrooms, with 21 ASVs shared between the two conditions. Alpha rarefaction analyses reached saturation for all sample groups, indicating that sequencing depth was sufficient to capture bacterial diversity (Supplementary Excel File 1). Alpha diversity differed significantly among the four experimental groups for Shannon diversity (*p* = 1.5 × 10^-^⁴), Faith’s phylogenetic diversity (*p* = 2.5 × 10^-^³), and observed features (*p* = 5.6 × 10^-^⁴) (Fig2A and Supplementary Excel File 1). In Farm X, symptomatic mushrooms exhibited significantly lower diversity across all three metrics than asymptomatic mushrooms (*p* < 0.05; Table S1). In Farm Y, disease status did not significantly affect Shannon diversity or observed features, whereas Faith’s phylogenetic diversity remained significantly different between disease states.

Beta-diversity analyses also revealed differences in bacterial community composition (Fig. 2B and Supplementary Excel File 1). Principal coordinates analysis (PCoA) based on Bray-Curtis dissimilarities showed partial separation of bacterial communities according to disease status and farm. Community composition differed significantly among groups (PERMANOVA, *pseudo-F* = 3.2762, *p* = 0.001). Pairwise comparisons were significant for most group combinations (*q* < 0.01; S1), with the exception of asymptomatic and symptomatic mushrooms from Farm Y (*pseudo-F* = 1.19, *q* = 0.302). Symptomatic samples also displayed greater dispersion within ordination space than asymptomatic samples.

**Figure 2.**
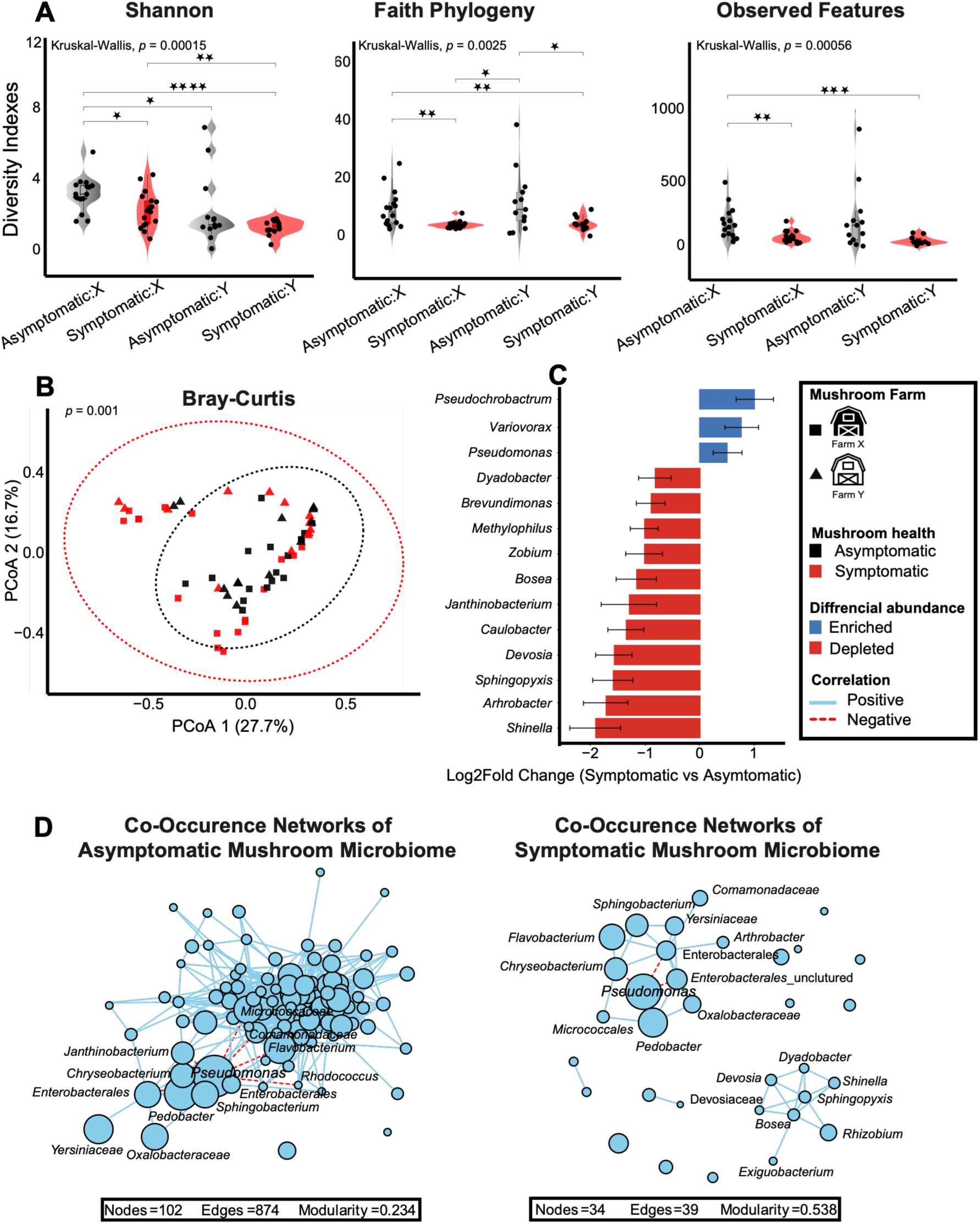
Community level restructuring of the mushroom bacteriome associated with bacterial blotch. (A) Alpha-diversity of bacterial communities based on Shannon diversity, Faith’s phylogenetic diversity, and observed features across asymptomatic and symptomatic mushrooms from Farms X and Y. Statistical significance was assessed using the Kruskal Wallis test. (B) Principal coordinates analysis (PCoA) based on Bray Curtis dissimilarity showing the separation of bacterial communities associated with asymptomatic and symptomatic mushrooms from Farms X and Y. Point shape indicates farm, and shading indicates mushroom health status. The percentages indicate the proportion of variation explained by each principal coordinate. PERMANOVA indicated a significant effect of mushroom health status on bacterial community composition (*p* = 0.001). (C) Differential abundance analysis was performed using ANCOM-BC to identify bacterial taxa significantly associated with mushroom health status. Log2 fold-change (log2FC) values represent the relative enrichment or depletion of bacterial taxa in symptomatic compared with asymptomatic mushrooms, with positive values indicating enrichment in symptomatic mushrooms and negative values indicating enrichment in asymptomatic mushrooms. Error bars represent standard errors of the estimated log2 fold-change. Taxa were considered significantly differentially abundant at *q* < 0.05. The 15 most enriched and 15 most depleted significant taxa are shown, with *Pseudomonas* highlighted separately (D) Co-occurrence networks of bacterial communities associated with asymptomatic and symptomatic mushrooms. Networks were constructed from significant pairwise Spearman correlations among bacterial genera detected in at least 20% of samples within each health-status group. Positive and negative associations are represented by distinct edge types. Node size indicates mean relative abundance, and network topology was evaluated based on connectivity and community structure.

Differential abundance analysis identified several bacterial genera associated with disease status (Fig. 2C and Supplementary Excel File 1). *Pseudochrobactrum* (log₂FC = 1.01, *q* = 0.012) and *Variovorax* (log₂FC = 0.77, *q* = 0.041) were significantly enriched in symptomatic mushrooms. In contrast, *Dyadobacter*, *Brevundimonas*, *Methylophilus*, *Rhizobium*, *Bosea*, *Janthinobacterium*, *Caulobacter*, *Devosia*, *Sphingopyxis*, *Arthrobacter* and *Shinella* were significant depleted (*q* < 0.05). Although *Pseudomonas* exhibited a positive log₂ fold change in symptomatic mushrooms (log₂FC = 0.52), the increase was not significant after multiple-testing correction (*q* = 0.135).

### Bacterial blotch is associated with reduced microbiome network complexity

Co-occurrence network analysis revealed marked differences in bacterial community organization between asymptomatic and symptomatic mushroom (Fig. 2D and Supplementary Excel File 1). The asymptomatic network comprised 102 nodes and 874 edges, with network density of 0.170, whereas the symptomatic network consisted only 34 nodes and 39 edges, resulting in substantially reduced connectivity (density = 0.070). In contrast, modularity increased from 0.234 in asymptomatic communities to 0.533 in symptomatic communities. The interaction profile of *Pseudomonas* also differed between disease states. In asymptomatic mushrooms, *Pseudomonas* exhibited nine significant negative associations with other bacterial taxa, including *Flavobacterium, Pedobacter, Sphingobacterium, Rhodococcus, Chryseobacterium,* and *Janthinobacterium* together with members of the Micrococcaceae and Comamonadaceae (SD1). In symptomatic mushrooms, only four significant negative associations were detected. Overall, symptomatic communities exhibited lower connectivity and fewer interactions involving *Pseudomonas*.

### Functional profiles of the disease-associated *Pseudomonas* communities

To investigate the metabolic potential of disease-associated bacterial communities, pathway abundances were compared between asymptomatic and symptomatic mushrooms using shotgun metagenomic data (Fig. 3). Both disease states contained many of the same dominant *Pseudomonas* taxa, including *P. gingeri*, *Pseudomonas* sp. NS1_2017, the *P. fluorescens* group, *P. agarici*, and *P. yamanorum*. Across taxa, the most abundant pathways were associated with fatty acid metabolism, carbon metabolism, amino acid biosynthesis, and nucleotide biosynthesis. Peptidoglycan maturation (PWY0-1586) represented the most abundant pathway and was associated with all major *Pseudomonas* taxa. Similarly, the O-antigen biosynthesis pathway (OANTIGEN-PWY) was widely distributed among dominant *Pseudomonas* species. *Pseudomonas* populations shared broadly similar functional profiles despite differences in taxonomic composition.

**Figure 3.**
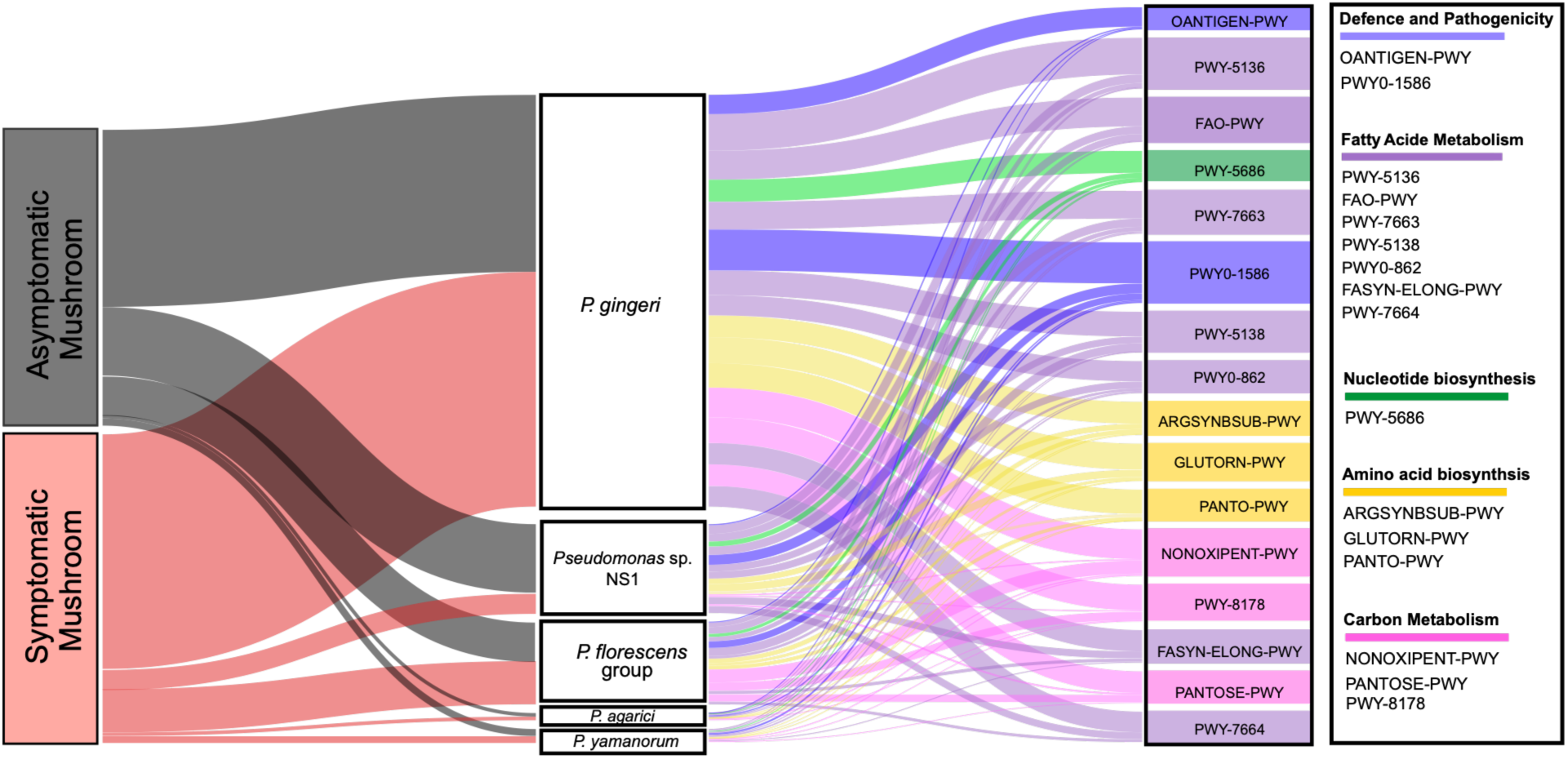
Pseudomonas-associated functional pathways across symptomatic and asymptomatic mushrooms. Sankey diagram showing the distribution of metabolic pathways across mushroom health status (left), individual pathogenic Pseudomonas taxa (middle), and their associated metabolic pathways (right). Flow widths and vertical block heights are proportional to the summed pathway abundance across samples. The 15 most abundant pathways were visualized. The far-right color bars indicate five major functional categories: (i) Defense and pathogenicity, Fatty acid metabolism, Nucleotide biosynthesis, Amino acid biosynthesis and Carbon metabolism. The differences in flow patterns and widths illustrate variation in the relative functional potential of pathogenic Pseudomonas communities associated with symptomatic versus asymptomatic mushrooms.

### Directional succession of casing microbiomes during mushroom development

To characterize temporal changes in the casing microbiome, bacterial communities were analyzed across the early, pinning, and harvest stages of mushroom production. Amplicon sequencing of 18 casing samples generated 14,663 bacterial ASVs with an average sequencing depth of 111,897 reads per sample. Shotgun metagenomic sequencing of nine representative casing samples from Farm X generated 170.1 million paired-end reads, of which 166.4 million high-quality reads were retained after quality filter (Supplementary Excel File 2)

Genus-level analyses revealed pronounced successional shifts in bacterial community composition throughout mushrooms development (Fig 4.A and Supplementary Excel File 2). Early-stage casing at both farms was dominated by *Exiguobacterium* and *Leuconostoc.* As development progressed, these taxa declined, whereas *Flavobacterium* and *Pedobacter* increased in abundance. In Farm X, *Pseudomonas* increased from approximately 2% in early-stage casing to 15% during pinning and 51% at harvest. A similar but less pronounced trend was observed in Farm Y, where *Pseudomonas* increased from approximately 1% to 3% and 9% across the same developmental stages.

**Figure 4.**
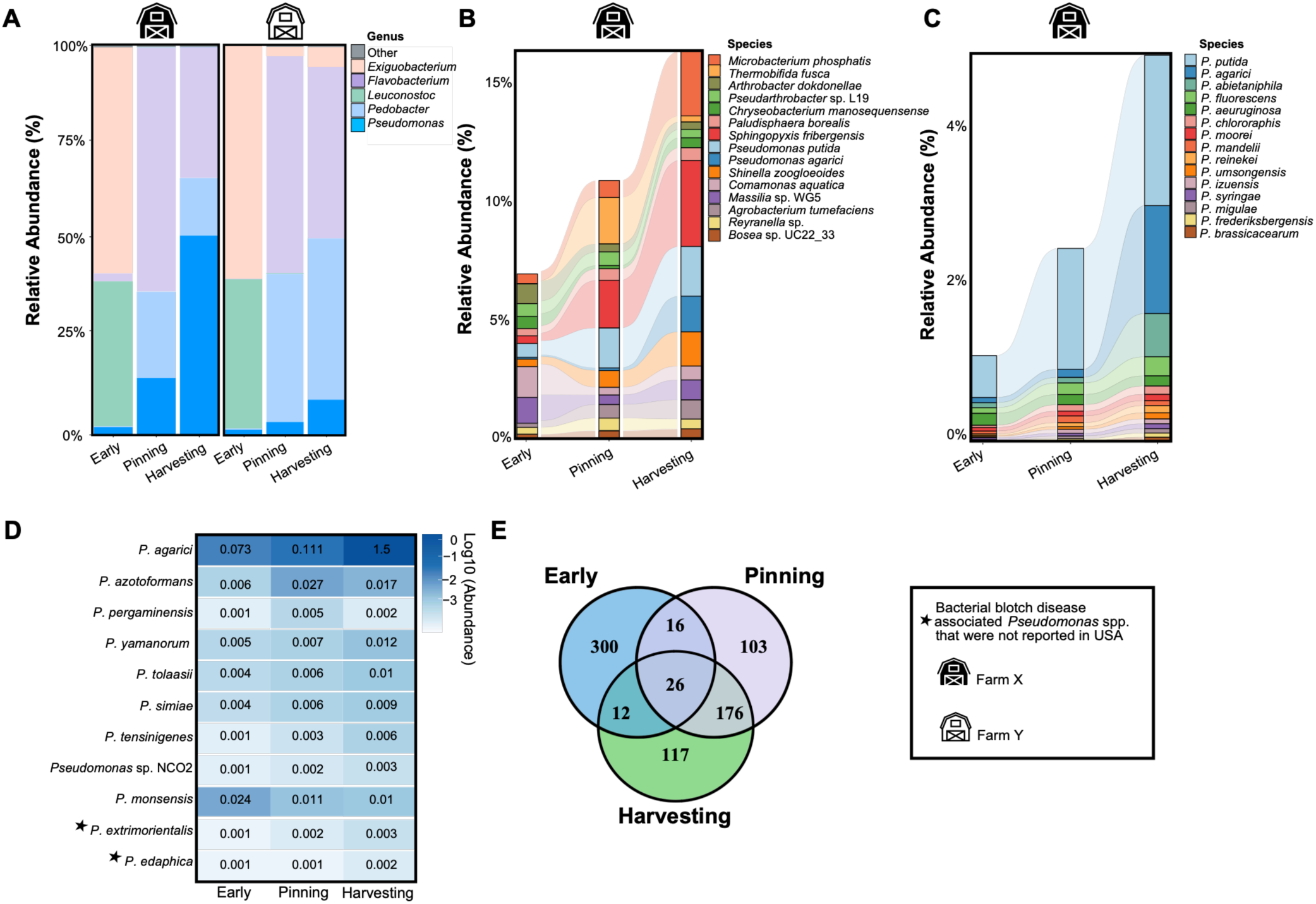
Temporal dynamics and diversity of bacterial blotch-associated *Pseudomonas* in the mushroom casing layer. (A) Genus-level relative abundance of the five most abundant bacterial genera in casing material collected at early, pinning, and harvesting stages from Farm X and Farm Y. The category “Other” represents all remaining bacterial genera. (C) Species-level relative abundance of the 15 most abundant bacterial taxa in casing material collected at early, pinning, and harvesting stages from Farm X. (D) Relative abundance of the 15 most abundant *Pseudomonas* species detected in casing material across early, pinning, and harvesting stages from Farm X. (E) Heatmap showing the relative abundance of bacterial blotch-associated pathogenic *Pseudomonas* species in casing material across early, pinning, and harvesting stages from Farm X. Representative symptoms associated with each *Pseudomonas* species are shown in the right panel, based on symptoms observed in isolates previously characterized by Mudiyanselage et al. (2026). Values represent mean log10-transformed abundance. (F) Venn diagram showing the numbers of bacterial ASVs shared among and exclusively detected at the early, pinning, and harvesting stages of mushroom development. * indicate *Pseudomonas* species associated with bacterial blotch disease that have not previously been reported in the United States. Farm labels indicate the sampling locations (Farm X and Farm Y).

Species-level metagenomic analysis revealed parallel shifts in bacterial community composition (Fig. 4B and Supplementary Excel File 2). Early-stage casing was dominated by *Massilia* sp. WG5 and *Comamonas aquatica*, whereas *Sphingopyxis fribergensis*, *Thermobifida fusca*, and *Pseudomonas putida* increased during pinning. Harvest-stage casing was characterized by greater representation of *S. fribergensis*, *Microbacterium phosphatis*, *P. putida*, *P. agarici*, and *Shinella zoogloeoides*. The composition of the *Pseudomonas* community also changed during casing development (Fig. 4C). Early-stage communities were dominated by *P. putida*, whereas later stages exhibited increased abundance and diversity of *Pseudomonas* species. By harvest, *P. putida* remained dominant, while *P. agarici* emerged as the second most abundant taxon. Screening casing metagenomes for bacterial blotch-associated *Pseudomonas* species identified eleven taxa throughout casing development (Fig. 4D). These included *P. tolaasii*, *P. agarici*, *P. yamanorum*, *P. simiae*, *P. azotoformans*, *P. pergaminensis*, *P. monsensis*, *P. tensinigenes*, *P. extremorientalis* and *P. edaphica* and *Pseudomonas sp.* NC02. All taxa occurred at relatively low abundances throughout casing development.

### Casing succession is characterized by extensive community restructuring

Core microbiome analysis revealed distinct bacterial assemblages associated with each developmental stage (Fig. 4E). Early-stage casing contained the highest proportion of unique core ASVs (40%), followed by harvest (15.6%) and pinning (13.7%). A total of 176 core ASVs (23.5%) were shared exclusively between the early-pinning and harvesting stages, whereas only 16 (2.1%) and 12 (1.6%) ASVs were shared exclusively between the early-pinning and early harvesting stages, respectively. Only 26 ASVs (3.4%) were shared across all three developmental stages.

Rarefaction curves reached saturation for all groups (Supplementary Excel File 2). Alpha diversity metrics, including Shannon diversity, Faith’s phylogenetic diversity, and observed ASVs, did not differ significantly among developmental stages (Kruskal-Wallis test, *P* > 0.05; Fig. 5A). In contrast, beta diversity analyses revealed significant shifts in community composition across casing development (Fig. 5B and Supplementary Excel File 2). The first two principal coordinates explained 57.6% of the total variation, with PC1 and PC2 accounting for 39.3% and 18.3%, respectively. Samples from the pinning and harvesting stages largely overlapped, whereas early-stage samples formed a distinct cluster separated from both the pinning and harvesting samples. Community composition differed significantly among developmental stages (*pseudo-F* = 6.33, *P*= 0.001), with significant differences between the early and pinning stages and between the early and harvesting stages. No significant differences were observed between the pinning and harvesting stages (*pseudo-F* = 1.70, *q* = 0.103; S2).

**Figure 5.**
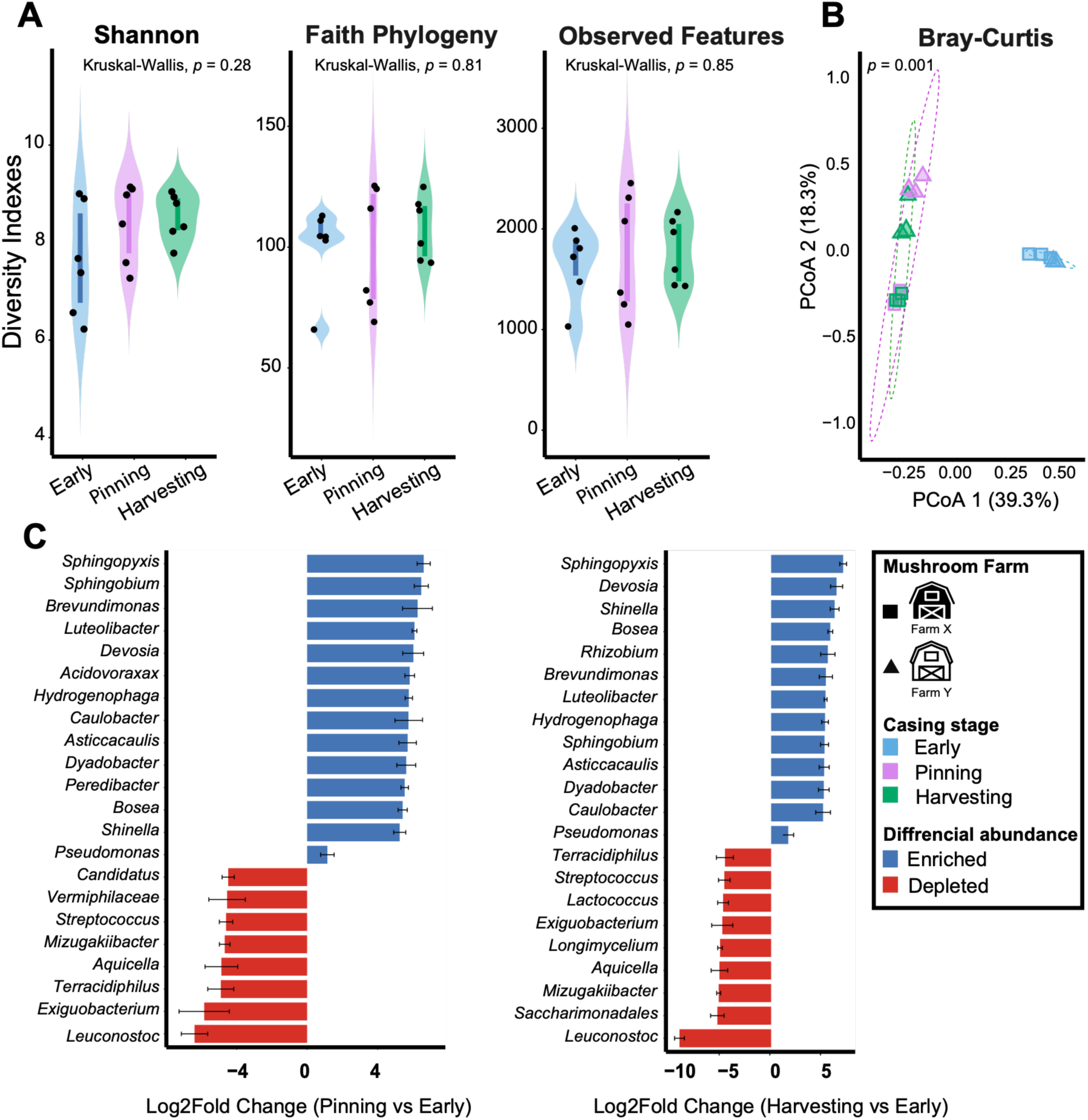
Community-level succession of mushroom casing bacteriome across developmental stages. (A) Alpha diversity of bacterial communities based on Shannon diversity, Faith’s phylogenetic diversity, and observed features across early, pinning, and harvesting stages at Farms X and Y. Statistical significance was assessed using the Kruskal Wallis test. (B) Principal coordinates analysis (PCoA) based on Bray Curtis dissimilarity showing the separation of bacterial communities across casing developmental stages at Farms X and Y. Point shape indicates farm, and shading indicates casing developmental stage. Percentages indicate the proportion of variation explained by each principal coordinate. PERMANOVA indicated a significant effect of casing developmental stage on bacterial community composition (*p* = 0.001). (C) Differential abundance analysis using ANCOM-BC to identify bacterial taxa significantly associated with casing developmental stages. Log2 fold-change (log2FC) values represent the relative enrichment or depletion of bacterial taxa in pinning versus early stages and harvesting versus early stages, with positive values indicating enrichment in the later developmental stage relative to the early stage and negative values indicating depletion. Error bars represent standard errors of the estimated log2 fold-change. Taxa were considered significantly differentially abundant at *q* < 0.05. The 15 most enriched and 15 most depleted significant taxa are shown, with *Pseudomonas* highlighted separately.

Differential abundance analysis identified further demonstrated extensive taxonomic turnover during casing succession (Fig. 5C and Supplementary Excel File 2). Genera including *Sphingopyxis*, *Sphingobium*, *Brevundimonas*, *Luteolibacter*, *Devosia*, *Hydrogenophaga*, *Caulobacter*, *Asticcacaulis*, *Dyadobacter*, *Bosea*, and *Shinella* increased significantly from the pinning stage through harvest. In contrast, *Leuconostoc*, *Exiguobacterium*, *Terracidiphilus*, *Aquicella*, *Mizugakiibacter*, and *Streptococcus* declined progressively throughout development. Although *Pseudomonas* increased in abundance during both pinning (log₂FC = 1.15) and harvest (log₂FC = 1.69) relative to the early casing, these changes were not statistically significant. Together, these findings demonstrate that casing microbiome undergo pronounced directional succession during mushroom production, characterized by major shifts in community composition and taxonomic replacement despite relatively stable levels of overall diversity.

## Discussion

Bacterial Blotch of cultivated white button mushroom has traditionally been viewed through the framework of individual bacterial pathogens, particularly *Pseudomonas tolaasii* and *P. “gingeri”* [7]. A growing diversity of *Pseudomonas* lineages, including *P. agarici*, *P. “reactans”*, *P. yamanorum*, *Pseudomonas* sp. NC02, *P. azotoformans*, *P. pergaminensis*, *P. monsensis*, *P. tensinigenes*, *P. edaphica*, *P. salomonii*, *P. extremorientalis*, *P. simiae*, *Pseudomonas* sp. Irchel 3A7, and *Pseudomonas* sp. REP124, has been associated with blotch symptoms [5,11]. The repeated detection of multiple *Pseudomonas* lineages together with diverse non-*Pseudomonas* taxa suggests that disease expression may be influenced by broader microbial community dynamics than by a single causal bacterium.

By integrating 16S rRNA gene amplicon sequencing, shotgun metagenomics, and temporal profiling of bacterial communities, our study demonstrates that bacterial blotch is associated with the reorganization of a *Pseudomonas*-dominated microbiome spanning both mushroom tissues and the casing layer. Rather than observing a transition from healthy mushrooms to communities dominated by a single pathogen, we identified changes in bacterial community composition, interaction structure, and microbial succession. These findings support an ecological model in which disease-associated populations emerge within an evolving microbial community, linking bacterial blotch development to the broader casing-mushroom microbiome continuum.

Bacterial blotch symptoms are generally expressed during later stages of mushroom development or post-harvest handling [35]. Consequently, the widespread dominance of *Pseudomonas* in both symptomatic and asymptomatic mushrooms provide important context for interpreting disease development (Fig. 1B). Across both farms researched in this study, *Pseudomonas* remained the predominant bacterial genus regardless the mushroom health. In Farm X, *Pseudomonas* represented approximately 77% and 78% of the bacterial community in symptomatic and asymptomatic mushrooms, respectively. This similarity is particularly notable given the substantially higher disease incidence at Farm X (∼30%) than at Farm Y (∼2%), with Farm X samples collected during the second flushing cycle, when disease was already present. These observations indicate that bacterial blotch cannot be explained simply by expansion of the *Pseudomonas* population. Instead, disease expression appears to occur within an already *Pseudomonas*-dominated community. This finding is consistent with growing evidence from plant and animal microbiome studies showing that disease states often arise through shifts in community composition and interactions rather than through overwhelming dominance of a single organism. Similar processes may operate in mushroom-associated microbiomes, where disease development could depend on factors such as pathogen establishment timing, host physiological status, cropping stage, environmental conditions, and interactions among co-occurring microbial populations [36,37]. Several bacteria genera, including *Pedobacter*, *Flavobacterium*, and *Chryseobacterium,* were also detected in both symptomatic and asymptomatic mushrooms, suggesting that these taxa are consistent members of the mushroom bacteriome, consistent with previous observations [38]. Notably, members of *Pedobacter* and *Sphingobacterium* detected on mushroom caps have previously been reported to detoxify tolaasin and suppress *P. tolaasii* under experimental conditions [39,40]. However, their presence in both asymptomatic and symptomatic mushrooms suggests that their contribution to disease may depend on strain identity, abundance, environmental conditions, and interactions with other community members. Species-level metagenomic analyses revealed contrasting shifts among dominant *Pseudomonas* lineages during disease development. *Pseudomonas* sp. Ost2, subsequently assigned by average nucleotide identity (ANI) analysis to the *P. “gingeri”* species complex (Supplementary Method), increased in relative abundance from 26.8% in asymptomatic to 42.5% in symptomatic mushrooms, whereas well-characterized blotch-associated taxa, including *P. tolaasii* and *P. azotoformans*, decreased in relative abundance (Fig. 1C). These opposing abundance patterns indicate that disease-associated changes within *Pseudomonas* reflect lineage-specific redistribution rather than a generalized expansion of blotch-associated taxa.

Changes involving non-*Pseudomonas* taxa reinforce this interpretation. *Ewingella americana* has previously been associated with internal stipe necrosis of cultivated mushrooms [41,42] and has been reported from mushroom production systems in the worldwide [41,43,44,45,46]. To our knowledge, *E. americana* association with cultivated mushrooms has not previously been documented in the United States. In the present study, *E. americana* decreased from approximately 2.6% in asymptomatic to ∼1.0% in symptomatic mushrooms, whereas *Mycetocola* sp. increased from 0.87% to 4.08% (Fig. 1C; Additional file 1). Interestingly, previous culture-based studies reported inhibitory activity of *E. americana* against *Mycetocola* sp. [41,38], and the contrasting abundance patterns observed here are consistent with a potential ecological relationship between these taxa, although this interaction remains to be experimentally tested. *Mycetocola* is particularly notable because its members exhibit diverse and potentially contrasting interactions with *Pseudomonas* spp aswell. *Mycetocola tolaasinivorans* and *M. lacteus* were originally isolated from mushroom fruiting bodies and shown to detoxify tolaasin and suppress *P. tolaasii in vitro* [40]. Conversely, other *Mycetocola* lineages have been associated with bacterial brown-pit disease of *A. bisporus* [12], suggesting that the ecological role of *Mycetocola* may depend on strain-level characteristics and microbial context. In this study, we further detected *Pseudomonas extremorientalis* and *Pseudomonas edaphica*, previously reported from blotch-associated communities outside the United States (Fig. 1E) [8,11]. Both taxa were detected in symptomatic and asymptomatic mushrooms, although at relatively low abundance.

Community-level analyses consistently indicated that symptomatic and asymptomatic mushrooms represent alternative configurations of a largely shared microbiome. Although symptomatic and asymptomatic mushrooms shared a substantial core bacteriome (∼42.9%), both groups also contained condition-associated ASVs (Fig. 1F). Moreover, alpha diversity patterns varied between farms (Fig. 2A). Farm X exhibited significant reductions in alpha diversity in symptomatic mushrooms, whereas Farm Y showed comparatively limited changes in richness and evenness. Such variation likely reflects differences in environmental conditions, crop management, disease pressure, and resident microbial communities among production systems [39,35]. Similarly, beta-diversity analyses revealed significant community differences without complete microbiome turnover (Fig. 2B). The greater stochastic dispersion observed among symptomatic mushrooms further indicates increased heterogeneity in community composition during disease, potentially reflecting alternative trajectories of colonization and disease progression among individual fruiting bodies. Importantly, *Pseudomonas* spp. itself was not significantly enriched in symptomatic mushrooms compared to the asymptomatic mushrooms (Fig. 2C). Instead, several less abundant genera, including *Pseudochrobactrum* and *Variovorax*, showed enrichment in symptomatic mushrooms, highlighting that disease-associated restructuring may extends beyond the dominant taxon and may involve lower-abundance community members as well.

The strongest evidence for disease-associated microbiome disruption emerged from co-occurrence network analyses (Fig. 2D). Symptomatic mushrooms exhibited dramatically reduced network complexity, with fewer nodes, fewer edges, and greater modularity than asymptomatic mushrooms. Similar reductions in network connectivity have been reported in dysbiotic microbiomes across diverse plant, animal, and environmental systems [47,48]. The decline in interactions involving *Pseudomonas*, together with the loss of associations with taxa such as *Pedobacter*, *Sphingobacterium, Flavobacterium*, and *Janthinobacterium*, suggests that disease development is accompanied by fragmentation of ecological interactions within the mushroom microbiome. Several of these taxa have previously been reported to exhibit potential antagonistic interactions with *Pseudomonas* [49,40,38]. Although co-occurrence networks cannot demonstrate causal interactions, they provide evidence that bacterial blotch is associated with reduced community connectivity and altered ecological organization.

Together, these taxonomic and network-level changes indicate substantial restructuring of the mushroom microbiome during bacterial blotch. We next examined whether this restructuring was accompanied by shifts in the functional potential of dominant *Pseudomonas* populations. Despite substantial taxonomic turnover, these populations exhibited broadly conserved functional profiles, with pathways involved in central carbon and fatty acid metabolism, amino acid and nucleotide biosynthesis, and cell-envelope biosynthesis consistently represented across dominant taxa (Fig. 3).The prominence of the peptidoglycan maturation pathway and broad representation of O-antigen biosynthesis among major *Pseudomonas* taxa are consistent with the fundamental importance of cell-envelope maintenance for bacterial growth, adaptation and defense within host-associated environments. Peptidoglycan provides structural support and is closely linked to the assembly and function of cell-envelope-associated structures, including systems involved in motility and environmental adaptation [50,51,52]. In damaged mushroom tissues, bacteria may encounter fluctuating moisture levels, oxidative compounds, altered oxygen availability, and host-derived metabolites such as tyrosinase generated during browning reactions [6]. Preservation of cell-envelope-associated functions may therefore support persistence under changing environmental conditions. The broad distribution O-antigens can also influence bacterial surface properties, interactions with the host, and resistance to external stresses [53]. However, metagenomic profiles reflect genetic potential rather than gene expression and therefore do not demonstrate active pathogenic functions.

One of the most significant findings of this study is the linkage between bacterial blotch-associated communities in mushroom tissues and microbial succession within the casing layer. The casing is essential for *Agaricus bisporus* production, providing both the physical environment required for fructification and a reservoir of microorganisms that influence mushroom development [14]. Previous studies have demonstrated that fluorescent *Pseudomonas*species, including *P. putida*and *P. poae*, contribute to fruit-body initiation and development [54,55,56]. Thus, the casing microbiome represents an integral component of mushroom biology rather than simply a growth substrate. Our temporal analyses revealed strong directional succession within casing bacterial communities. Early-stage casing was dominated by *Exiguobacterium* and *Leuconostoc*, whereas *Flavobacterium*, *Pedobacter*, and *Pseudomonas* progressively increased during pinning and harvest (Fig. 4A), consistent with previously reported successional patterns in mushroom production systems [48]. Particularly noteworthy was the progressive increase in *Pseudomonas* throughout casing development (Fig. 4A), rising from approximately 2% in early-stage casing to 51% at harvest in Farm X. Species-level analyses revealed increasing representation of *P. putida*, along with blotch pathogen *P. agarici* (Fig. 4B). In addition, blotch-associated *Pseudomonas* taxa previously detected in mushroom cap tissues were also detected at low abundance across casing stages (Fig. 4D). Together, these findings suggest that the casing layer may serve as a potential reservoir of diverse *Pseudomonas* populations with potentially contrasting ecological roles, ranging from beneficial interactions with the mushroom to association with disease.

All those changes in the casing microbial community occurred without significant differences in alpha diversity, indicating that succession primarily involved replacement and redistribution of taxa rather than a simple gain or loss of bacterial diversity (Fig. 5A). The strong separation of early-stage casing from pinning and harvest communities further indicates that the major ecological transition occurred relatively early in mushroom development (Fig. 5B). In contrast, the substantial overlap between pinning and harvest communities indicates that the casing microbiome becomes comparatively stable after this transition. Such directional restructuring may reflect changes in nutrient availability, fungal development, moisture, pH, oxygen availability, and host-derived compounds as the crop progresses [17].

## Conclusion

Our findings establish bacterial blotch disease in cultivated white button mushroom as a microbiome-associated disease shaped by microbial succession, community restructuring, and host development across the casing mushroom continuum. Rather than being characterized by the proliferation of a single pathogen, disease was associated with pronounced restructuring of bacterial communities, including species-level shifts within *Pseudomonas*, changes in non-*Pseudomonas* taxa, and reduced microbial network connectivity. The progressive enrichment of *Pseudomonas* during casing development, together with the detection of multiple blotch-associated lineages in casing material, further identifies the casing layer as a potential reservoir and ecological filter for bacteria that subsequently associate with developing fruiting bodies.

These findings extend the conventional pathogen-centered view of bacterial blotch by demonstrating that disease-associated bacteria occur within a dynamic microbial ecosystem in which their abundance and ecological context change throughout mushroom development. Importantly, the contrasting responses of closely related *Pseudomonas* lineages and other bacterial taxa indicate that disease emergence cannot be inferred from pathogen presence or abundance alone. Instead, the ecological state of the surrounding community may influence which bacterial populations persist, proliferate, or become associated with disease.

From a disease-management perspective, the casing microbiome represents a potential intervention point. Understanding how casing composition, cultivation practices, and microbial succession shape the establishment of beneficial versus disease-associated populations could provide new opportunities for disease prediction and management while preserving microbial functions essential for mushroom production. Future studies integrating longitudinal sampling with strain-resolved metagenomics, culture-based interaction assays, and experimental manipulation of casing communities will be essential for determining the mechanisms underlying these community transitions.

## Data availability

The 16S rRNA gene amplicon and shotgun metagenomic sequencing reads are available in the NCBI Sequence Read Archive (SRA) under BioProjects PRJNA1523947 and PRJNA1523863, respectively.

## Author Contributions

S.J.M. obtained the funding source; S.D.M., S.J.M., and R.G. designed the research; S.D.M. and S.J.M performed experiments; S.D.M. wrote the manuscript; R.G., and S.J.M reviewed the manuscript.

## Acknowledgments

The authors also acknowledge Dr. Yuru Chang for the assistance with the mushroom maceration process.

